# The transcriptional regulator JAZ8 interacts with the C2 protein from geminiviruses and limits the viral infection in *Arabidopsis*

**DOI:** 10.1101/2022.08.11.503596

**Authors:** Tabata Rosas-Diaz, Pepe Cana-Quijada, Mengshi Wu, Hui Du, Gemma Fernandez-Barbero, Alberto P. Macho, Roberto Solano, Araceli G. Castillo, Xiao-Wei Wang, Rosa Lozano-Duran, Eduardo R. Bejarano

## Abstract

Jasmonates (JAs) are phytohormones that finely regulate critical biological processes, including plant development and defense. JASMONATE ZIM-DOMAIN (JAZ) proteins are crucial keeping JA-responsive genes in a repressed state. In the presence of JA-Ile, JAZ repressors are ubiquitinated and targeted for degradation by the ubiquitin/proteasome system, allowing the activation of the downstream transcription factors and, consequently, the activation of JA-responsive genes. A growing body of evidence has shown that JA signalling is crucial in defending against plant viruses and their insect vectors. Here, we describe the interaction of C2 proteins from two geminiviruses from the genus Begomovirus, tomato yellow curl Sardinia virus (TYLCSaV) and tomato yellow leaf curl virus (TYLCV), with the transcriptional repressor JAZ8 from *Arabidopsis thaliana* and its closest orthologue in tomato, SlJAZ9. Both JAZ and C2 proteins colocalize in the nucleus, forming discrete nuclear speckles. Overexpression of *JAZ8* did not lead to altered responses to TYLCV infection; however, knock-down of *JAZ8* favours the geminiviral infection in plants. Low levels of *JAZ8* likely affect the viral infection specifically since *JAZ8*-silenced plants do not display developmental phenotypes nor present differences in their interaction with the viral insect vector. Our results show that JAZ8 interacts with geminiviral C2 proteins and exerts an anti-geminiviral effect.

## INTRODUCTION

Phytohormones are chemical messengers essential in establishing signalling networks to regulate plant growth and stress-related responses (Gimenez-Ibanez et al., 2017), jasmonic acid (JA) and its methyl ester (MeJA) and isoleucine conjugate (JA-Ile) are formed from fatty acids and are collectively known as jasmonates (JAs); jasmonates control critical biological processes in plants, including fertility, seedling development, response to wounding, and the growth-defense trade-off (Gimenez-Ibanez et al., 2017; Howe et al., 2018; Ruan et al., 2019). When encountering stress, such as those that results from insect feeding or the attack by necrotrophic fungi, eudicots rapidly biosynthesize JA and its bioactive form, JA-Ile (or dn-OPDA in bryophytes; Monte et al., 2018), to trigger signal transduction events, which ultimately lead to the onset of plant defense (reviewed in Campos et al., 2014; Yan and Xie, 2015; Howe et al., 2018). In basal conditions, JAs are maintained at low levels, and JA-mediated transcriptional responses are kept repressed by JASMONATE ZIM-DOMAIN (JAZ) proteins (Wasternack and Song, 2017). JAZ proteins repress the activity of crucial JA-inducible transcription factors (TFs) such as the basic helix–loop–helix (bHLH) MYC2 and its homologs MYC3/4/5. These TFs, in turn, regulate the expression of a large proportion of JA-responsive genes, including the expression of *JAZ* genes, to attenuate JA responses as part of a feedback loop pathway (Chico et al., 2008; Howe et al., 2018; Zander et al., 2020). In response to specific stresses, there is an increase in the levels of bioactive JA-Ile. Importantly, this compound binds the receptor protein CORONATINE INSENSITIVE1 (COI1) and acts as molecular glue, facilitating the interaction between the JAZ repressors and COI1, the adaptor subunit of the E3 ubiquitin ligase complex SCF(SKP1/Cullin1/F-box)^COI1^ (Xie et al., et al., 1998; Fonseca et al., 2009; Chini et al., 2007). This hormone-dependent interaction leads to JAZs degradation through the ubiquitin/26S proteasome pathway, allowing the de-repression of the downstream TFs and, consequently, the activation of JA-responsive genes (Howe et al., 2018).

There are thirteen JAZ proteins (JAZ1-13) in *Arabidopsis thaliana* (hereafter referred to as *Arabidopsis*). The members of the JAZ family exhibit high sequence variability but generally possess two common domains: ZIM, which includes the conserved TIFY (TIFF/YXG) motif, and Jas (Chini et al., 2016; Guo et al., 2018; Thireault et al., 2015; Garrido-Bigotes et al., 2019). Within the Jas domain, a degron signal can be found, which is responsible for the degradation of JAZs in the presence of JA–Ile and may play a role in nuclear localization (Chini et al., 2007; Thines et al., 2007; Melotto et al., 2008; Grunewald et al., 2009).

The absence of solid phenotypes in most *jaz* single mutants and the analysis of *Arabidopsis* mutants defective in multiple *JAZ* indicated a relatively high level of redundancy among JAZ family members (Campos et al., 2016; Chini et al., 2016; Major et al., 2017; Guo et al., 2018), limited by tissue-specific expression patterns of some *JAZ* genes.

Some JAZ proteins, including *Arabidopsis* group IV (JAZ7, JAZ8, and JAZ13, and alternative splice variants of JAZ10), are recalcitrant to COI1 interaction since they harbour a noncanonical degron and therefore present a greater stability and repressor activity in a jasmonate-stimulated context (Chung et al., 2009; Shyu et al., 2012; Thireault et al., 2015; Howe et al., 2018). JAZ8, when ectopically expressed in *Arabidopsis*, represses JA-regulated defense responses and senescence (Shyu et al., 2012; Jiang et al., 2014; Chen et al., 2021). Interestingly, JAZ8 has been described as a component of the so-called JJW (JAV1-JAZ8-WRKY51) complex, which finely controls JA biosynthesis genes to defend against insect attack (Yan et al., 2018). Besides, Chen and collaborators have determined that *JAZ8* overexpression repressed plant defense against *Botrytis cinerea* by interacting with the transcription factor WRKY75, which positively regulates JA-mediated plant defense against necrotrophic fungal pathogens. JAZ8 also interacts with VirE3 from *Agrobacterium tumefaciens* to attenuate root tumorigenesis by antagonistically modulating (the salicylic acid/JA)-mediated plant defense signalling (Li et al., 2021b).

Geminiviruses represent a large family of insect-transmitted plant viruses with circular single-stranded DNA genomes (ssDNA, ∼2.7–5.2 kb) packaged within geminate particles. Begomovirus, the largest genus within the *Geminiviridae* family, comprises viruses transmitted by whiteflies that can have either mono-or bipartite-genomes traditionally considered to encode six to eight proteins (Zerbini et al., 2017; Fiallo-Olivé et al., 2021). Despite their limiting coding capacity, the genome-encoded information is sufficient to complete all processes required for infection, such as viral replication, movement, and suppression or evasion of plant defense mechanisms (reviewed in (Fondong, 2013; Hanley-Bowdoin et al., 2013; Ramesh et al., 2017; Aguilar et al., 2020). It has been reported that geminiviruses have evolved strategies to subvert JA signalling, which plays an important role in the tripartite interaction between plants, geminiviruses, and their insect vectors (Sun et al., 2017; Wu and Ye, 2020; Pan et al., 2021). Repression of the JA pathway or JA-responsive genes has been demonstrated in *Arabidopsis*, tomato, and tobacco plants infected with begomoviruses as well as in transgenic plants expressing geminiviral pathogenicity factors like the C2 or the βC1 proteins (Ascencio-Ibáñez et al., 2008; Yang et al., 2008; Lozano-Durán et al., 2011a; Li et al., 2014; Rosas-Díaz et al., 2016; Shi et al., 2019; Li et al., 2019b; Guerrero et al., 2020). Interestingly, exogenous application of MeJA negatively impacts geminiviral infection (Lozano-Durán et al., 2011a; Chakraborty and Basak, 2019).

C2 represents the multifunctionality of geminiviral proteins (Guerrero et al., 2020). This small viral protein (∼15 KDa) localizes mainly in the nucleus and could act as a transcriptional factor for viral genes (Sunter and Bisaro, 1992). It is likewise able to trigger the transcription of host genes (Trinks et al., 2005; Oh et al., 2006; Lozano-Durán et al., 2011a; Rosas-Díaz et al., 2016; Li et al., 2019a), and suppresses transcriptional and post-transcriptional gene silencing (Dong et al., 2003; Wang et al., 2003; Vanitharani et al., 2004; Wang et al., 2005; Buchmann et al., 2009; Luna et al., 2012). Transgenic expression of the C2 protein from the begomoviruses tomato yellow curl Sardinia virus (TYLCSaV) and tomato yellow leaf curl virus (TYLCV) or from members of the genera curtovirus such as beet curly top virus (BCTV) subverts ubiquitination of the COP9 signalosome complex (CSN), affecting cellular processes regulated by SCF complexes, including JA-signalling (Lozano-Durán et al., 2011a; Lozano-Durán et al., 2011b). However, transcriptomic analyses show that plants expressing C2 from TYLCSaV are only affected in specific JA-induced responses, implying that C2 alters JA-dependent gene regulation by additional mechanisms leading to specificity (Rosas-Díaz et al., 2016). Moreover, C2 from TYLCV interacts with a ubiquitin-related protein, RPS27A, compromising the degradation of JAZ1 and resulting in the inhibition of JA-mediated plant defense (Li et al., 2019a).

While the role of C2 in suppressing JA-triggered responses has been demonstrated for several geminiviruses, whether other plant proteins besides RPS27A interact with C2 and contribute to its specific effect on JA responses remains to be determined. In this study, we found that the C2 proteins from TYLCSaV and TYLCV interact with JAZ8 from *Arabidopsis* and its closest orthologue in tomato, SlJAZ9, colocalizing in the nucleus. *Arabidopsis* plants with low sensitivity to JA due to overexpression of *JAZ8* did not show altered responses to TYLCV infection; however, the lack of JAZ8 favours the viral infection. Finally, we observed that JAZ8 did not significantly contribute to the performance of the whitefly *Bemisia tabaci*, the insect vector for begomoviruses, in *Arabidopsis*. Our results show that JAZ8 interacts with C2 in the plant cell nucleus and suggest that JAZ8 exerts a direct anti-geminiviral effect.

## RESULTS

### Exogenous application of jasmonates hampers TYLCV infection in *Arabidopsis*

It was previously shown that exogenous MeJA (JA) treatment has a negative effect on the infection by BCTV in *Arabidopsis* (Lozano-Durán et al., 2011a). To determine if the application of JA also has an impact on the infection by begomoviruses, we inoculated JA- and mock-treated *Arabidopsis* plants with TYLCV. Four to five-week-old plants inoculated with TYLCV were treated with 50 μM JA or mock solution every two days starting at two days post-inoculation (dpi). The transcript accumulation of JA-responsive genes and the amount of viral DNA were determined at 16 and 21 dpi, respectively. The results show that the application of exogenous JA reduced the accumulation of viral DNA (Figure 1A). The efficacy of the JA treatments was confirmed by the induction of JA-responsive genes (*AtAOS, AtJAZ8*, and *AtJAZ10*) in infected and uninfected plants treated with JA (Figure 1B). Interestingly, while in the non-treated plants, the presence of TYLCV does not alter the transcript accumulation of *AtAOS* and *AtJAZ10* compared to non-infected plants, the expression of *AtJAZ8* is induced by the viral infection. However, the elevation in *JAZ8* transcripts triggered by TYLCV does not affect its induction by JA (Figure 1B).

**Figure 1.**
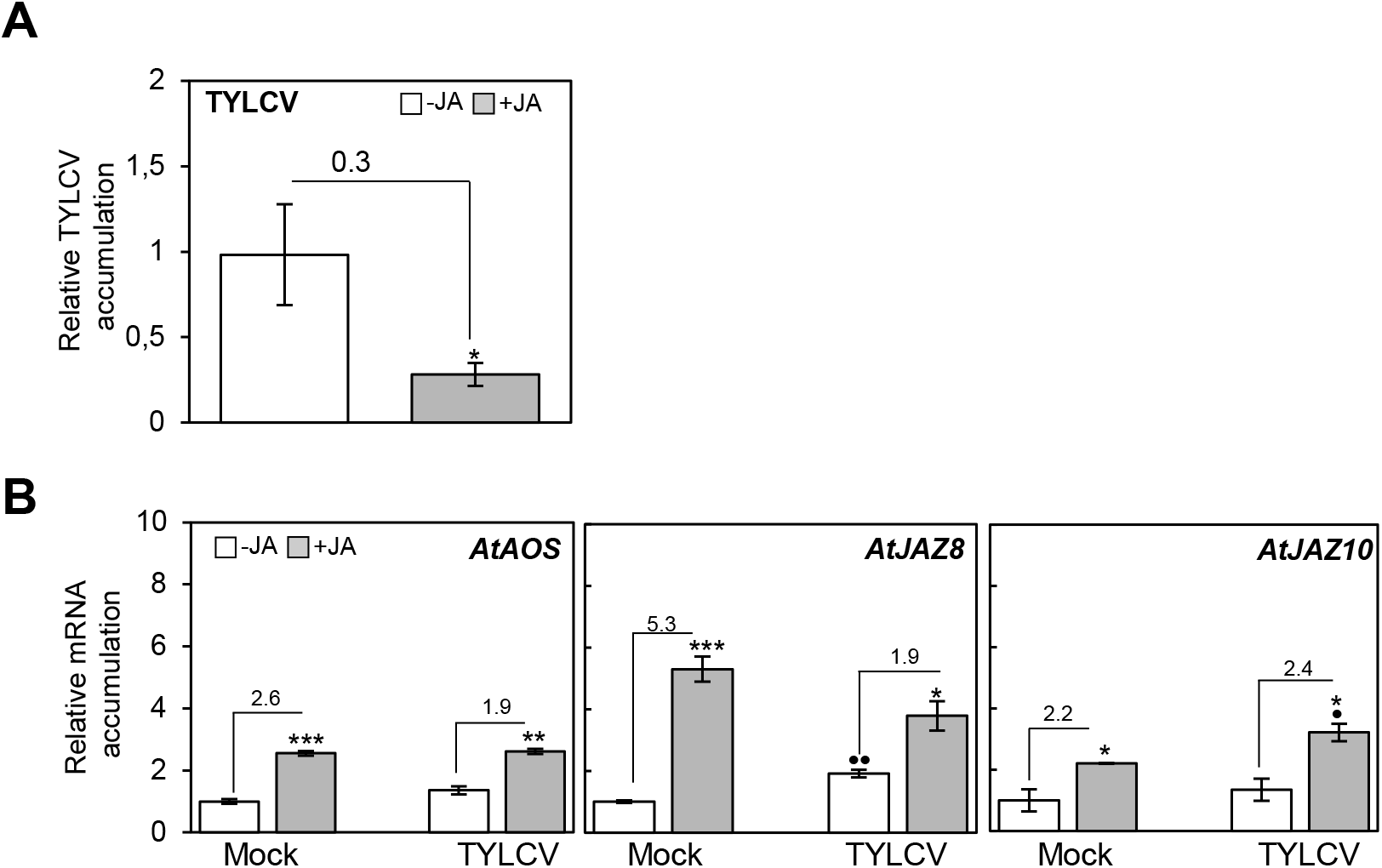
Exogenous JA application negatively impacts TYLCV infection, and geminivirus infection alters JA signalling in *Arabidopsis*. Four-to five-week-old *Arabidopsis* plants were agroinoculated with TYLCV/Mld and, at 2 dpi, sprayed every other day with 50 μM MeJA (+JA) or mock solution (-JA) (0.5% ethanol in water (v/v)). (**A**) Relative TYLCV DNA accumulation was determined by real-time PCR of total DNA extracted from whole plants at 21 dpi. Average values of twelve infected plants are represented. Bars represent standard error. The asterisk indicates a statistically significant difference between untreated and treated samples with *p-value < 0.05, according to a Student’s t-test. **(B)** Expression of JA-responsive genes upon exogenous treatments in *Arabidopsis* plants. Mock-(pBINX) and TYLCV-infected plants were treated with 50μM MeJA (+JA) or mock solution (-JA); JA responsive genes *AtAOS* (At542650), *AtJAZ8* (At1g30135), and *AtJAZ10* (At5g13220) were quantified by real-time RT-qPCR at 17 days post-treatment. The average values of the three plants are represented. Bars represent standard error. Asterisks (between –JA vs. +JA) or dots (pBIN vs. TYLCV) indicate a statistically significant difference compared to the relevant control (***, p-value < 0.005; **/··, p-value < 0.01; */·, p-value < 0.05), according to a Student’s t-test. Three independent experiments were performed in A and B with similar results; results from one representative replicate are shown.

### C2 proteins from TYLCV and TYLCSaV interact with the transcriptional repressor JAZ8 from *Arabidopsis* and its orthologue in tomato

JAZ proteins have been identified as targets of plant pathogen effectors, including viruses (Wu et al., 2017; Li et al., 2019a; Oblessuc et al., 2020; Yang et al., 2020). To further investigate the mechanism conferring specificity to the C2-mediated interference with JA responses (Rosas-Díaz et al., 2016), we analyzed if C2 was able to interact with JAZ proteins. Thus, we tested the interaction between C2-TS1-78, a truncated version of the C2 protein from TYLCSaV lacking the autoactivation domain (Lozano-Durán et al., 2011a), and eleven members of the JAZ family from *Arabidopsis* by yeast two-hybrid. The results showed that this viral protein interacts with AtJAZ8 (Figure 2A). To confirm this interaction *in planta*, we conducted co-immunoprecipitation (Co-IP, Figure 2B) and Förster resonance energy transfer-fluorescence lifetime imaging microscopy (FRET-FLIM, Figure 2C) assays using transient expression in *Nicotiana benthamiana*. In addition, to determine if this interaction is conserved, we included in the analyses C2 from another tomato-infecting begomovirus, TYLCV (C2-TY) (Figure S1), and the closest tomato orthologue of AtJAZ8, SlJAZ9 (Figure S2). For Co-IP, *N. benthamiana* leaves were co-infiltrated with constructs to express JAZ proteins (RFP-tagged at the C-terminus) and the geminiviral C2 proteins (GFP-tagged at the N-terminus). As a negative control, leaves were co-infiltrated with a construct to express free GFP. As shown in Figure 2B, both GFP-C2 proteins could associate with AtJAZ8-RFP and SlJAZ9-RFP, while free GFP did not. In the input, we observed one intense band corresponding to the predicted size of JAZ proteins fused to RFP (around 40 KDa) and a heavier but faint band (approximately 50 KDa) that could correspond to post-translational modifications by ubiquitin or SUMO, previously detected in JAZ proteins (Gough and Sadanandom, 2021). Notably, JAZ proteins co-immunoprecipitated with both C2 proteins showed these two bands with similar intensities.

**Figure 2.**
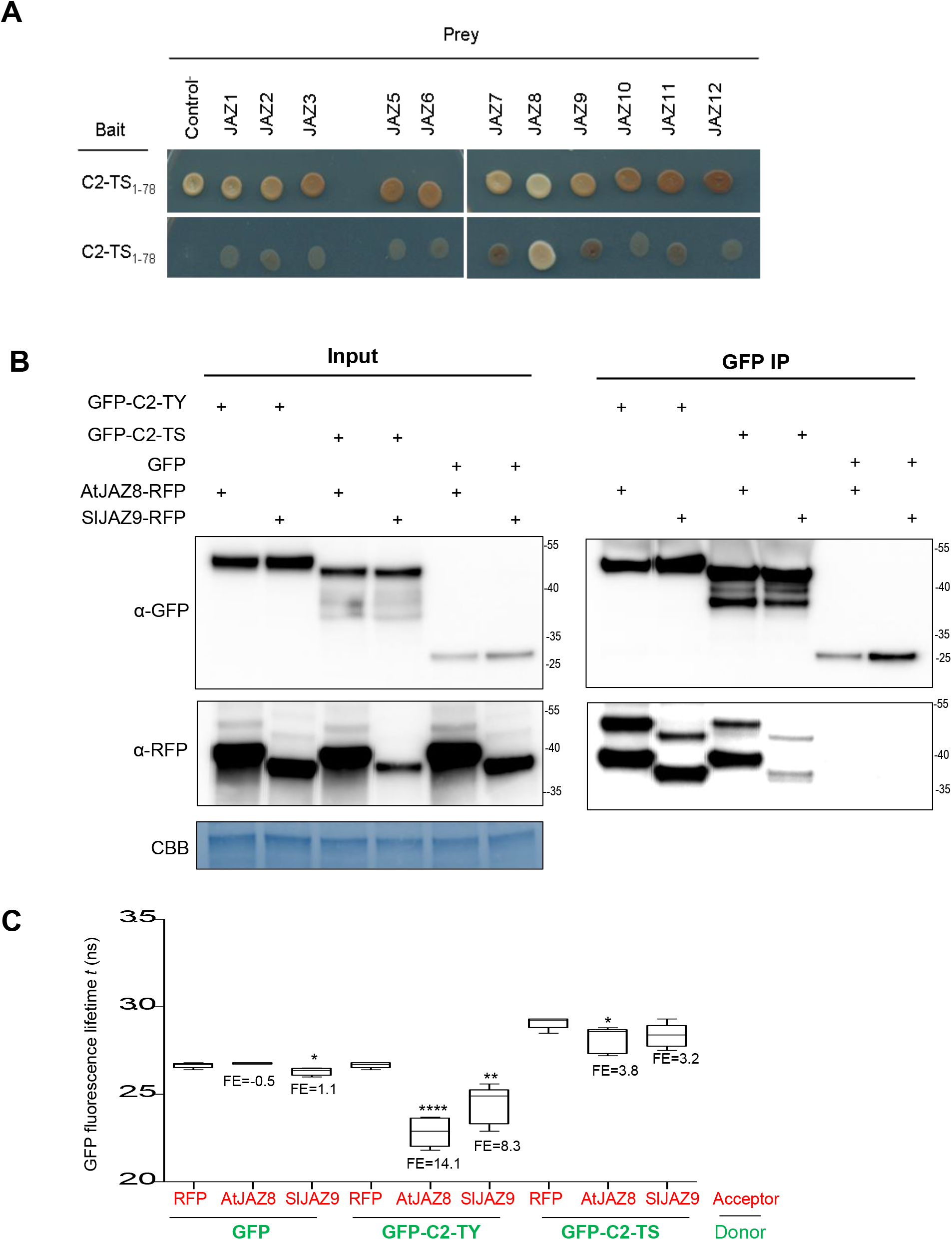
C2 from TYLCV and TYLCSaV interact with JAZ8 from *Arabidopsis* and JAZ9 from tomato. **(A)** Yeast two-hybrid with C2 from tomato yellow leaf curl Sardinia virus (C2-TS) and *Arabidopsis* JAZ proteins. Yeast cells co-transformed with pGBKT7-C2-TS(1-78) (bait) and pGADT7-JAZ (prey) were selected and subsequently grown on a medium lacking Leu and Trp (-2), as a co-transformation control, or on selective medium lacking Ade, His, Leu and Trp (-4) to test protein interactions. As a control, pGBTKT7-C2-TS was co-transformed with the pGADT7 vector. **(B)** Co-immunoprecipitation analysis of C2 from tomato yellow leaf curl virus (C2-TY) and tomato yellow leaf curl Sardinia virus (C2-TS) with AtJAZ8 and SlJAZ9 from *Arabidopsis* and tomato, respectively, upon transient co-expression in *N. benthamiana* leaves. Molecular weight is indicated. CBB: Comassie brilliant blue. **(C)** Interaction between 35S:GFP-C2-TY/-C2-TS (donors) and 35S:AtJAZ8/SlJAZ9-RFP (acceptors) by FRET-FLIM upon transient co-expression in *N. benthamiana* leaves. Free GFP is used as a negative control. FE: FRET efficiency. Asterisks indicate a statistically significant difference (****,p-value < 0.0001; *, p-value < 0.05), according to a Student’s t-test.

To evaluate if the viral proteins and the transcriptional repressors are located in the same cellular compartment and whether one alters the subcellular localization of the other, we transiently expressed them, alone or in combination, fused to fluorescent proteins in *N. benthamiana* leaves (Figure 3). We observed that both C2 proteins (GFP-tagged at the N-terminus) are localized in the nucleoplasm and excluded from the nucleolus. In addition, C2 from TYLCSaV (C2-TS) also accumulated at the cell periphery. Like the viral proteins, AtJAZ8 (RFP-tagged at the C-terminus) localizes in the nucleoplasm and is excluded from the nucleolus. SlJAZ9 also localizes in the nucleoplasm but shows a trend to accumulate surrounding the nucleolus. Strikingly, the sub-nuclear localization of AtJAZ8 and C2 proteins changes when they are co-expressed, with both colocalizing in discrete nuclear speckles (C2-TY) or forming aggregates all over the nucleoplasm (C2-TS). However, only C2 from TYLCV changed subnuclear localization when co-expressed with SlJAZ9 by positioning around the nucleolus. These results indicate that C2 from begomoviruses interacts with both AtJAZ8/SlJAZ9 in the nucleus.

**Figure 3.**
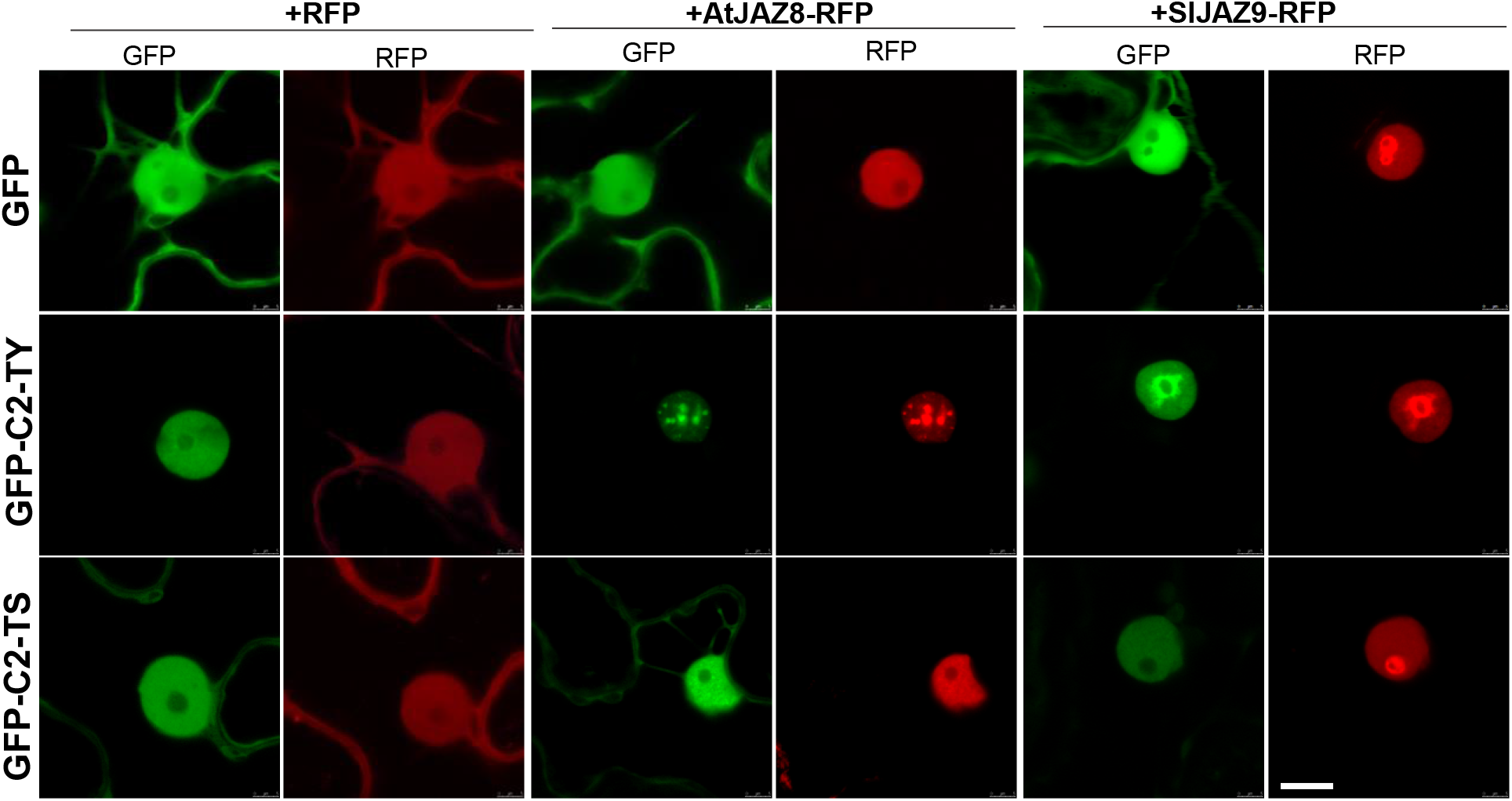
AtJAZ8 and SlJAZ9 co-localizes with C2 from TYLCV and TYLCSaV in the nucleus. Subcellular co-localization of 35S:GFP-C2-TY/C2-TS and 35S:AtJAZ8/SlJAZ9-RFP upon transient co-expression in *N. benthamiana* leaves at two days post-infiltration. GFP: GFP channel, RFP: RFP channel. Bar represents 10 μm.

### AtJAZ8 is not required for normal plant development

To study the relevance of AtJAZ8 in the repression of JA responses, we characterised *Arabidopsis* lines altered in *AtJAZ8* expression. To reach this aim, we employed a previously described over-expressing transgenic line, *35S:AtJAZ8-GUS* (*JAZ8*-OX, Shyu et al., 2012), that accumulates 20 times more *JAZ8* transcript than the wild type (WT) (Figure 4B), and generated *AtJAZ8* knock-down lines expressing the precursor of an artificial miRNA targeting the *AtJAZ8* transcript (amiRNA*JAZ8*; hereafter referred to as amjaz8 plants) (Figure 4A). Two amjaz8 homozygous lines (Line 19 and 20) showing a drastic reduction of the *AtJAZ8* transcript accumulation (between 87-97% compared to the WT; Figure 4B) were selected and used for further analysis. As previously reported by Shyu et al. (2012), the increase in *AtJAZ8* expression does not significantly affect plant morphology or size (Figure 4C). Similarly, no noticeable differences were observed in the transgenic plants showing lower accumulation of *AtJAZ8* either (Figure 4C).

**Figure 4.**
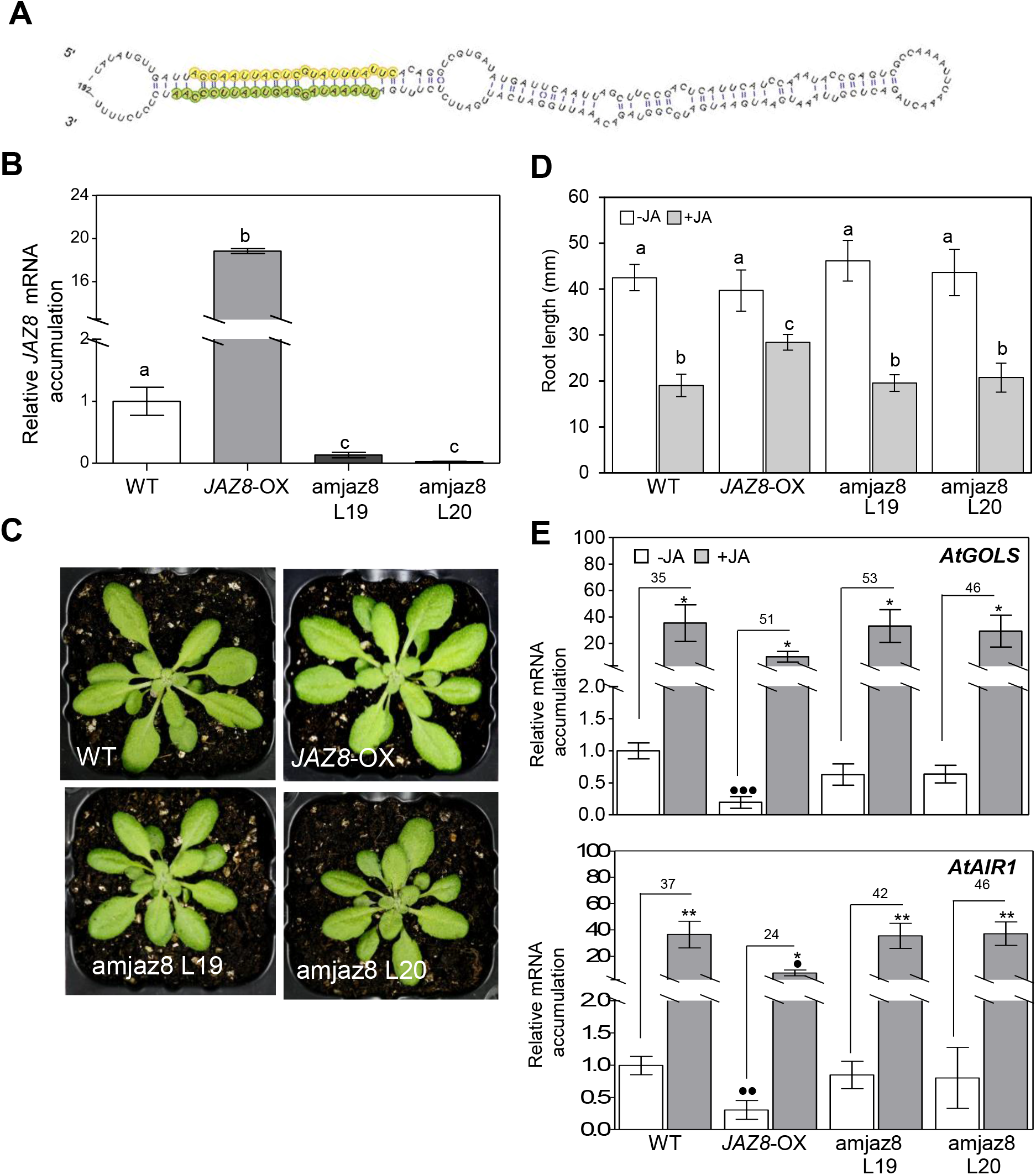
JAZ8 is not required for normal plant development (A). Generation of *Arabidopsis JAZ8*-knockdown line using artificial microRNAs. Secondary structure of the artificial micro RNA precursor targeting *JAZ8* (amjaz8). The mature amjaz8 and amjaz8* sequences are highlighted in green and yellow. Secondary structure was predicted using Mfold (Zuker, 2003) and visualized using varna (http://varna.lri.fr/). **(B)** Pictures of four-week-old *Arabidopsis* plants 35S:JAZ8 (*JAZ8*-OX) and two independent amjaz8 homozygous lines (L19, L20) compared with wild type (WT) plants. **(C)** Relative *JAZ8* mRNA levels in wild type (WT), 35S:*JAZ8* (*JAZ8*-OX) and amjaz8 seedlings. Total RNA was extracted and subjected to quantitative RT-PCR (RT-qPCR) analysis to measure the *JAZ8* mRNA levels normalized to *ACTIN 2*. Values are represented as the relative expression compared to WT. Bars represent the mean +/-SD for three different pools of 8-10 seedlings. Columns with the same letters are not significantly different (P=0.05) according to Dunnett’s multiple comparison test. Experiments were repeated twice with similar results; results from one representative experiment are shown. **(D)** Root growth inhibition assays in 35S:JAZ8 (*JAZ8*-OX), amjaz8, and control wild type (WT) *Arabidopsis* seedlings growing in 50 μM MeJA (+JA) or mock media (-JA) (0.5% ethanol in water (v/v)). **(E)** Relative expression levels of *AtGOLS* (At2g47180) and *AtAIR1* (At4g12550) genes in seedlings from (A), determined by real-time PCR. mRNA levels were normalized to *ACTIN. 2*. Values are represented as the relative expression compared to WT. Bars represent the mean +/-SD for three different pools of 8-10 seedlings. Asterisks (between –JA vs. +JA) or dots (–JA vs. -JA or +JA vs +JA) indicate a statistically significant difference compared to the relevant control (···, p-value < 0.005; **/··, p-value < 0.01; */·, p-value < 0.05), according to a Student’s t-test. t. Experiments were repeated twice with similar results; results from one representative experiment are shown.

Overexpression of *AtJAZ8* affects the sensitivity to JA (Shyu et al., 2012). To determine whether the reduction of *AtJAZ8* expression has a similar effect, we carried out a growth inhibition assay using amjaz8 plants. *JAZ8*-OX plants were used as control. The results showed that the reduction of *AtJAZ8* transcripts does not alter the root growth inhibition produced by exogenous JA application (Figure 4D).

Our previous work showed that the induction of several jasmonate-responsive genes was partially suppressed in transgenic plants expressing C2 from TYLCSaV (Rosas-Díaz et al., 2016). To determine whether changes in *AtJAZ8* expression affect the expression of two of those genes, *AtGOLS* and *AtAIR1*, we measured the transcript accumulation in the *JAZ8*-OX and amjaz8 lines. No significant changes in the expression were observed in the amjaz8 lines, either JA-treated or untreated. However, *JAZ8*-OX plants showed an apparent reduction in the expression of these marker genes in both treated and untreated plants, suggesting that overexpression of *AtJAZ8* mimics the effect of *C2* expression (Figure 4E).

Of note, when analysing an *AtJAZ8* line containing a T-DNA insertion in the gene promoter (*jaz8-1*), previously described as a *jaz8* knock-down line (Jiang et al., 2014) (Figure S3), we found that homozygous T-DNA plants over-accumulate *AtJAZ8* transcripts (Figure S3D), but their JA-induced inhibition of root growth is not altered (Figure S3E-F). Although we do not know the cause of the differences in the JA-response phenotype observed in both overexpressing lines (*JAZ8*-OX and *jaz8-1*), a dose-effect could be hypothesised, since *AtJAZ8* transcript accumulation is four times higher in the *JAZ8*-OX plants (Figure 4AB, S3D).

### JAZ8 limits the TYLCV infection in *Arabidopsis*

Under natural conditions, TYLCV is transmitted by the whitefly *Bemisia tabaci*. However, most of the viral infection experiments performed in laboratory conditions are carried out using *Agrobacterium*-mediated infection. Considering that alterations in the salicylic acid (SA) and jasmonate-mediated signalling pathways could affect the efficiency of *Agrobacterium*-mediated T-DNA transfer (Rosas-Díaz et al., 2017), we tested the competence of T-DNA transfer in WT, *JAZ8*-OX, and amjaz8 lines. Leaves were infiltrated with *Agrobacterium* containing a binary plasmid with a GUS-intron construct (Zipfel et al., 2006), which allows expression of the reporter in plants but not in bacteria. We included the *Arabidopsis NahG* transgenic plants as control, which have enhanced susceptibility to *Agrobacterium* transient transformation (Rosas-Díaz et al., 2017). As shown in Figure S4, in control WT leaves, only weak GUS staining was detectable at four dpi; *JAZ8*-OX and amjaz8 leaves showed similar GUS staining to WT plants, while *NahG* leaves exhibited stronger staining, as expected. This result reveals that neither overexpression nor lack of *JAZ8* altered the *Agrobacterium*-mediated transient transformation in *Arabidopsis*. Next, to evaluate the relevance of AtJAZ8 in the geminiviral infection, we agroinoculated *JAZ8*-OX and amjaz8 lines with TYLCV since TYLCSaV is unable to infect this plant species (Cañizares et al., 2015). As shown in Figure 5, the viral accumulation at 21 dpi in *JAZ8*-OX was similar to that in WT plants, but, strikingly, viral accumulation was significantly higher in both amjaz8 lines (Figure 5).

**Figure 5:**
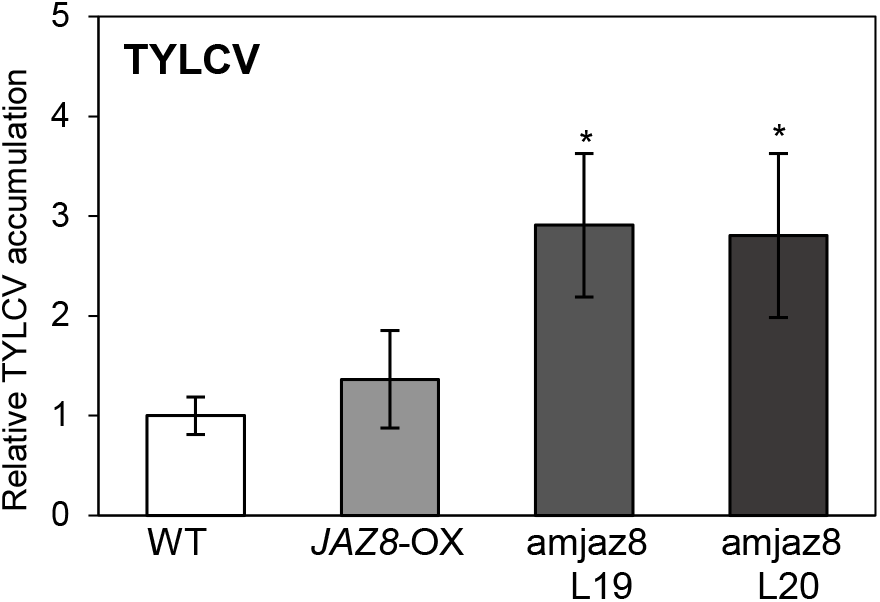
amjaz8 plants are more susceptible to TYLCV infection. Four to five-week-old wild type (WT), 35S:*JAZ8* (*JAZ8-*OX), and amjaz8 lines *Arabidopsis* plants were agroinoculated with TYLCV. The relative accumulation of viral DNA was determined by quantitative real-time PCR of total DNA extracted from whole plants at 21 days post-inoculation (dpi). Average values of 8-10 infected plants are represented. Bars represent standard error. Asterisks indicate statistically different samples from the control sample (*, p-value < 0.05) according to a Student’s t-test. Three independent experiments were performed with similar results; results from one representative replicate are shown.

Therefore, the lack of JAZ8 favours viral accumulation, suggesting that this transcriptional repressor limits the infection.

### JAZ8 does not affect the insect vector’s performance in the plant

The whitefly *B. tabaci* suppresses jasmonate-mediated defenses in *Arabidopsis* (van de Ven et al., 2000; Kempema et al., 2007; Zarate et al., 2007; Walling, 2008). Moreover, *Arabidopsis* and tomato plants impaired in JA-defenses show increased oviposition and enhanced development of the whitefly nymphs (Sanchez-Hernandez et al., 2006; Zarate et al., 2007; Zhang et al., 2013; Zhang et al., 2018). Since a plethora of viral proteins have emerged as manipulators of JA-mediated responses (Westwood et al., 2014), a scenario where the impairment of this pathway improves the performance of viral insect vectors, hence viral propagation, has been proposed (reviewed in Csorba et al., 2015; Zhang et al., 2018; Wu and Ye, 2020). To test whether JAZ8 also plays a role in the *Arabidopsis* response to whiteflies, we examined the effect of JAZ8 on insect performance, determining whitefly survival and fecundity on *JAZ8*-OX, amjaz8, and WT plants. Per each assay, ten adult whiteflies (5 males and 5 females) were collected and released on the *Arabidopsis* plants; seven days later, the survival of adults and the number of eggs on all three genotypes were quantified. Our tests showed that neither the enhanced accumulation nor depletion of the *AtJAZ8* transcript significantly affected the survival and fecundity of whiteflies (Figure 6).

**Figure 6.**
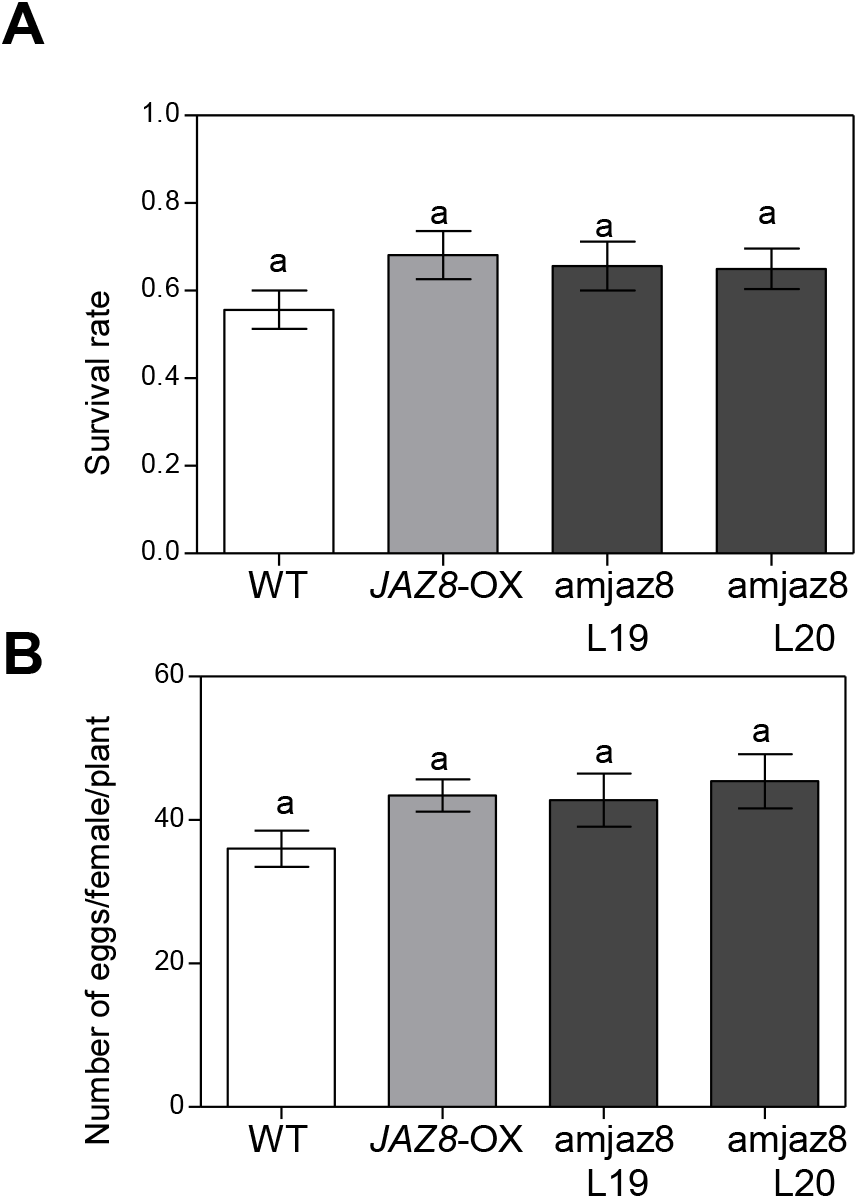
JAZ8 has no effect on the virus’ insect vector, the whitefly *Bemisia tabaci*. Performance of whitefly adults on the different *Arabidopsis* genotypes. Seven days after the release of whitefly adults, survival of adults **(A)** and the number of eggs per female (**B**) on wild type control (WT), 35S:*JAZ8* (*JAZ8*-OX) and amjaz8 were compared. Values are means± SE, n=16. Different letters denote significant differences (Student’s t-test, P<0.05).

## DISCUSSION

Besides its impact on the begomovirus vector fitness, jasmonate responses also positively affect the plant antiviral defense. Our results with jasmonate-treated plants infected by TYLCV confirmed that induction of jasmonate-response negatively affects viral accumulation (Lozano-Durán et al., 2011a; Jia et al., 2016; Chakraborty and Basak, 2019). The relevance of this hormonal response against geminiviruses is further supported by the identification of viral proteins able to hinder it: at least two geminiviral proteins have evolved mechanisms to reduce the intensity of the jasmonate response. In begomoviruses associated with betasatellite, a satellite-encoded protein, βC1, interferes with this pathway at several levels: (i) reducing the JAZ degradation by impairing the integrity of SCF^COI1^ complex through its interaction with SKP1 (Jia et al., 2016); (ii) suppressing of JA-responsive genes involved in defense against insects by interaction with the ASYMMETRIC LEAVES 1 (AS1) (Yang et al., 2008); or (iii) blocking the activation of terpene synthase genes by binding the basic helix-loop-helix transcription factor MYC2 (Li et al., 2014). In geminiviruses not associated with betasatellite, the virus-encoded C2 protein may play a similar role in limiting the jasmonate-response. C2 impacts ubiquitination at least by two mechanisms: (i) obstructing the correct assembly and disassembly of the SCF E3 ligases, which depend on the activity of the CSN complex, and, therefore, must affect the function of all hormone responses dependent on SCF function (Lozano-Durán et al., 2011a); (ii) indirectly compromising the degradation of JAZ1, and therefore impairing the JAZ1-dependent plant defense responses (Li et al., 2019a).

The JA-signalling pathway regulates multiple plant processes, including development, growth, and defense. The F-box protein COI1 perceives jasmonates, but the complexity of the response is orchestrated by the degradation of JAZ repressors and the release of numerous TFs, including MYC2 and its homologues. Although JAZ proteins were initially assumed to be functionally redundant, differential expression patterns, protein interactions, sensitivity to COI1-dependent degradation, and altered JA responses in several loss- and gain-of-function mutants suggest that they also have specific functions (Goossens et al., 2016; Chini et al., 2016). The functional diversity of the JAZ proteins and the fact that other pathogen-derived proteins from RNA viruses (Wang et al., 2014; Wu et al., 2017; Yang et al., 2020; Li et al., 2021a), bacteria (reviewed in Schreiber et al., 2021), or oomycetes (Mukhtar et al., 2011) interact with JAZs to suppress JA signalling and plant defenses, brought us to explore those transcriptional regulators as potential additional targets of the multifunctional C2 protein. Although C2 could indirectly compromise the degradation of AtJAZ1 (Li et al., 2019a), a direct interaction of C2 with the JAZ proteins has not been reported.

We show here that C2 interacts with one member of the group IV of JAZ proteins, AtJAZ8 from *Arabidopsis*, and its closest orthologue in tomato, SlJAZ9 (Figure 2). The fact that viral accumulation is higher in *AtJAZ8*-silenced plants indicates that the presence of JAZ8 limits the infection. Notably, overexpression of *JAZ8* does not alter viral accumulation, suggesting that JAZ8 is not rate-limiting for its antiviral effect or that other rate-limiting factors are involved in this process. Like all JAZ proteins, JAZ8 modulates the expression of downstream genes by interacting with many transcription factors, including two members of the NAC (NAM, ATAF1/2, and CUC2) family of proteins (ANAC062 and ANAC091) (Altmann et al., 2020) that function as regulators of defense-related genes (Bian et al., 2021). ANAC091, which is involved in hypersensitive response-mediated resistance to turnip crinkle virus (TCV) (Ren et al., 2000), functions through transcriptional activation to promote a basal level of resistance in the plant, while ANAC062 induces a group of *PR* genes, including *PR1, PR2*, and *PR5*, in an SA-independent manner. In accordance with this mechanism, overexpression of an active NAC062 form exhibited enhanced disease resistance to *Pseudomonas syringae* (Seo et al., 2010). Those NAC genes may participate in the plant response to geminivirus infection: infection with cabbage leaf curl virus induced the expression of *ANAC062* in *Arabidopsis* (Ascencio-Ibáñez et al., 2008), while TYLCV infection of tomato plants altered the expression of a large number of annotated *NAC* genes, including induction of *ANAC062/091* tomato homologues (*SlNAC25* and *SlNAC55*) (unpublished data and Huang et al., 2017).

Considering these results, we propose a model to explain the impact of the JA-induced response on the geminivirus infection (Figure 7). Jasmonate-dependent genes involved in plant defense are mainly controlled by the JAZ proteins sensitive to degradation mediated by the COI1-JA complex. In contrast, JAZ8, which hampers the geminiviral infection, cannot associate strongly with COI1 in the presence of JA-Ile. When plants are infected with geminiviruses, C2 interferes with the jasmonate-dependent response to the viral propagation through impairing the function of SCF complexes and so affecting all COI1-dependent responses and interacting with JAZ8 protein. The result is a level of viral accumulation in the plant that we consider a wild-type infection. Interestingly, this defect on the viral infection by JAZ8 seems specific to the JA-mediated response to pathogen infection, because amjaz8 plants were not affected in JA development responses or whitefly’s fitness (Figures 4 and 6). The fact that *JAZ8* silencing favours the viral infection indicates that *JAZ8* is negatively impacting the virus’ ability to infect the plant. Whether this negative effect of JAZ8 depends on the expression of *JAZ8*-regulated plant genes or it is a consequence of a direct inhibitory effect of JAZ8 over C2, which gets released when the former is silenced, remains to be clarified.

**Figure 7.**
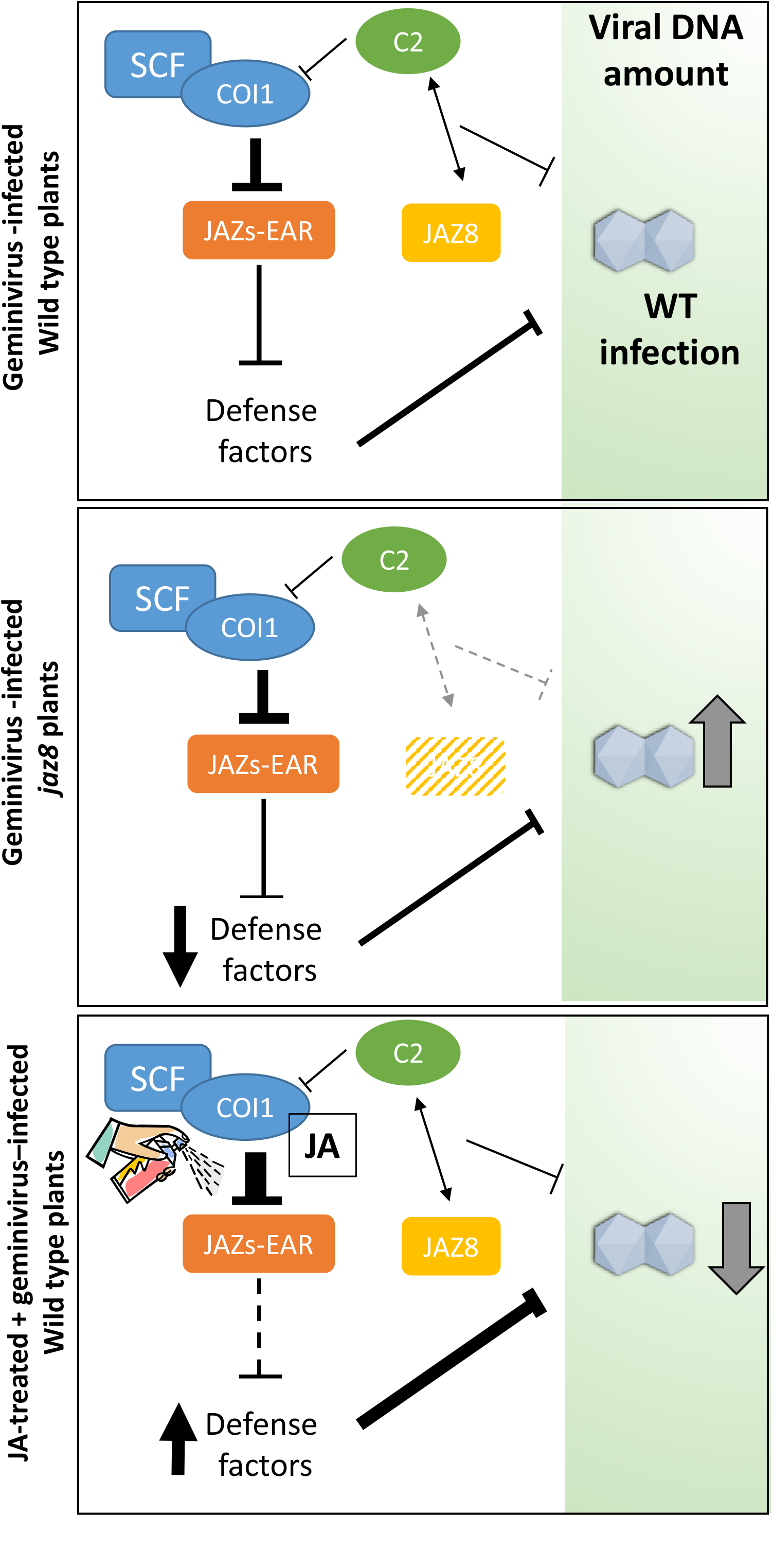
Tentative model that may explain the impact of the JA-induced response in the geminivirus infection. **(A)** When plants are infected with geminiviruses, C2 interferes with the jasmonate-dependent response to the viral propagation through impairing the function of SCF complexes and so affecting all COI1-dependent responses and by interacting with the JAZ8 protein. The result is a level of viral accumulation in the plant that we consider a wild-type infection. (B) The fact that *JAZ8* silencing favours the viral infection indicates that *JAZ8* is negatively impacting the viral ability to infect the plant. Whether this negative effect of JAZ8 depends on the expression of *JAZ8*-regulated plant genes or it is a consequence of a direct inhibitory effect of JAZ8 over C2 which gets released when the former is silenced, remains to be clarified. Finally, (C) when JA is applied, there is a burst of expression of defense-related genes controlled by JA-sensitive JAZ proteins, while the expression of the JAZ8-dependent genes remains unaffected. This induction of defense-related genes negatively affects the viral infection, which explains the reduction in viral accumulation observed in the JA-treated plants.

When JA is applied, there is a burst of expression of defense-related genes controlled by JA-sensitive JAZ proteins, while the expression of the JAZ8-dependent genes remains unaffected. This induction of defense-related genes might negatively affect the viral infection, which would explain the reduction in viral accumulation observed in the JA-treated plants.

In summary, the data presented here, revealing that JAZ8 has an antiviral function and that C2 and AtJAZ8/SlJAZ9 proteins physically interact, offer additional support to the relevance of JA-dependent responses in plant defense against geminiviruses. However, the fact that SA-dependent responses also participate in setting up a defense program against geminiviruses (Li et al., 2019b; Medina-Puche et al., 2020) draws a complex scenario played by these two hormone-mediated pathways setting up the plant defense against those viruses.

## MATERIAL AND METHODS

### Microorganisms and general methods

Manipulations of *Escherichia coli* and *Saccharomyces cerevisiae* strains and nucleic acids were performed according to standard methods (Ausubel et al., 1989; Sambrook and Russell, 2001). *Agrobacterium tumefaciens* (*Agrobacterium*) GV3101 strain was used for the agroinfiltration in *Nicotiana benthamiana* and *Arabidopsis thaliana* (*Arabidopsis*) and for geminiviral infections in *Arabidopsis. S. cerevisiae* strain pJ696 (*MATa, trp1-901, leu2-3,112, ura3-52, his3-200, gal4*Δ, *gal80*Δ, *GAL2-ADE2, LYS2::GAL1-HIS3, met2::GAL7-lacZ*), a derivative of PJ69-4A (James et al., 1996), was used for the yeast two-hybrid experiments.

Plant DNA extraction using CTAB method was performed as described in (Lukowitz et al., 2000). Plant RNA extraction was isolated using TRIzol reagent (Invitrogen).

### Plant materials and growth conditions

*Arabidopsis* plants accession Columbia (Col-0) wild type (WT), and mutant or transgenic derivatives were grown in growth chambers with 8 h light: 16 h dark cycles at 21°C. The T-DNA insertion *jaz8-1* line mutant (WiscDsLox255G12) was provided by the Nottingham *Arabidopsis* Stock Centre (NASC; http://www.Arabidopsis.info) and transgenic *35S:JAZ8* is described in Shyu et al., 2012.

Root growth inhibition assay in *Arabidopsis* seedlings was performed as described in Rosas-Díaz et al., 2016.

*N. benthamiana* plants were grown in soil at 24ºC in long-day conditions 16 h light: 8 h dark photoperiod.

### Transient expression assays

Transient co-expression was performed in 3-to 4-week-old *N. benthamiana* leaves through *Agrobacterium* infiltration (OD600 = 0.5) as described in Rosas-Díaz et al., 2018. For co-IP, FRET-FLIM, and subcellular co-localization assays, clones expressing GFP-C2 were co-infiltrated with clones to express JAZ-RFP (see “Plasmids and cloning”). Samples were taken two days after infiltration.

*Agrobacterium*-mediated expression in *Arabidopsis* was performed as described in Rosas-Díaz et al., 2017.

### Plasmids and cloning

Plasmids are summarized in supplementary table 1. Vectors from the pGWB series were described in Nakagawa et al., 2007. pB7RWG2.0 was described in Karimi et al., 2002. pBIN19-35S::GUS was kindly provided by Dr. Zipfel, The Sainsbury Laboratory, Norwich, UK.

*AtJAZ8* (At1g30135) and *SlJAZ9* (Solyc08g036640.2) genes were amplified by PCR (without stop codon) and cloned into pENTR/D-TOPO vector (Invitrogen) using the primers listed in Supplemental Table 2 generating TOPO-*AtJAZ8* and TOPO-*SlJAZ9*, respectively.

### Yeast two-hybrid assay

Yeast two-hybrid assays using the fragment from 1-78 aa of C2 of tomato yellow leaf curl Sardinia virus (C2-TS) and JAZ proteins from *Arabidopsis* were performed as described in Chini et al., 2009.

### Protein extraction and Co-immunoprecipitation

Two days after infiltration, 0.5 g of infiltrated *N. benthamiana* leaves were harvested. Protein extraction, Co-immunoprecipitation (Co-IP), and western blot were performed as previously described in Rosas-Díaz et al., 2018. The antibodies used are as follows: anti-GFP (Abiocode M0802-3a), anti-RFP (Chromotek 5F8), anti-Rabbit IgG (Sigma A0545), and anti-Mouse IgG (Sigma A2554).

### FRET-FLIM imaging

Two days after infiltration, *N. benthamiana* leaf discs co-expressing the proteins of interest were used to perform FRET-FLIM experiments. GFP was used as a donor protein, whereas the proteins fused to RFP were used as acceptor proteins. FRET-FLIM analysis was performed as described previously (Rosas-Díaz et al., 2018).

### Geminivirus infection assays

TYLCV infections of *Arabidopsis* plants were performed by agroinoculation (Lozano-Durán et al., 2011a). Plants were agroinoculated with pBINX’ (mock) or TYLCV/Mld (AF071228). Samples were taken at 21 dpi. Viral DNA accumulation was quantified by quantitative real-time PCR with primers listed in supplementary Table 2. MeJA treatments for the geminiviral infection experiments were carried out as done by (Lozano-Durán et al., 2011a)

### Quantitative Real-Time PCR (RT-qPCR)

cDNA preparation and RT-qPCR were performed as previously described by Rosas-Díaz et al., 2016. Primers are listed in Supplemental Table 2.

### Construction of *AtJAZ8* artificial miRNA and generation of transgenic plants

To generate the 21nt artificial microRNA (amiRNA) against *AtJAZ8* (amjaz8), we used Web MicoRNA Designer 3 (WMD3) (Ossowski et al., 2008). Using WMD3 we obtained the sequence of the presumably best amiRNA candidate and the primers used to amplify this sequence (supplemental Table 2). Using PCRs and overlapping PCRs as indicated in WMD3 protocol, we cloned the 21nt sequence of interest into the miR319 precursor backbone. The PCR fragment was gel-purified using Wizard SV gel and PCR Clean-up System (Promega), ligated into pGEM-T (Promega, USA), and checked by sequencing. Finally, this construct was subcloned into pBINX’ plasmid using *Sal*I (Takara, Japan) and *BamH*I (Takara, Japan). pBINX’-am*jaz8* or the empty vector (EV) were transformed into *Agrobacterium. Arabidopsis* plants were transformed using the floral dipping method (Clough and Bent, 1998). Transgenic seeds were selected on ½ standard Murashige and Skoog medium containing Kanamycin (50 mg/l). T3 antibiotic-resistant transformed plants were verified by sequencing. Repression of *JAZ8* was verified on T3 antibiotic-resistant transformed plants by RT-PCR and qPCR, using primers listed in supplemental Table 2.

### *Bemisia tabaci* bioassay

The whitefly *B. tabaci* MEAM1 (mtCOI GenBank accession: GQ332577) was maintained on 6-7th true-leaf stage cotton (*Gossypium hirsutum*) in a climate-controlled chamber with 14h light: 10h dark cycles at 27±1°C, 70±10% RH. Three days post-emergence, ten adult whiteflies (5 males and 5 females) were collected, and then released into a modified Lock&Lock Box with a host *Arabidopsis* placed inside. Sixteen replicates were conducted. Seven days later, the survival of adults was recorded, and all the eggs laid were counted to assess host plant suitability.

### *Arabidopsis* leaves GUS staining

GUS staining was performed according to the protocol previously described by Rosas-Díaz et al., 2017.

**Supplementary table 1.**
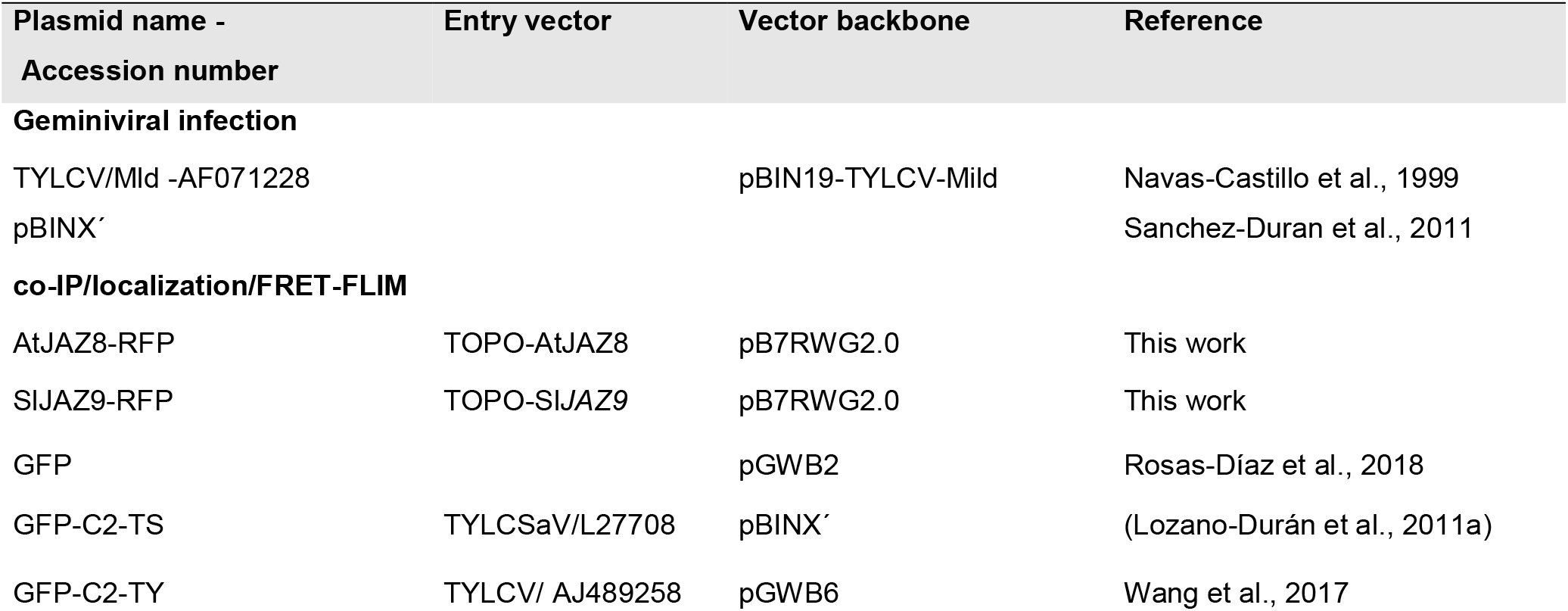
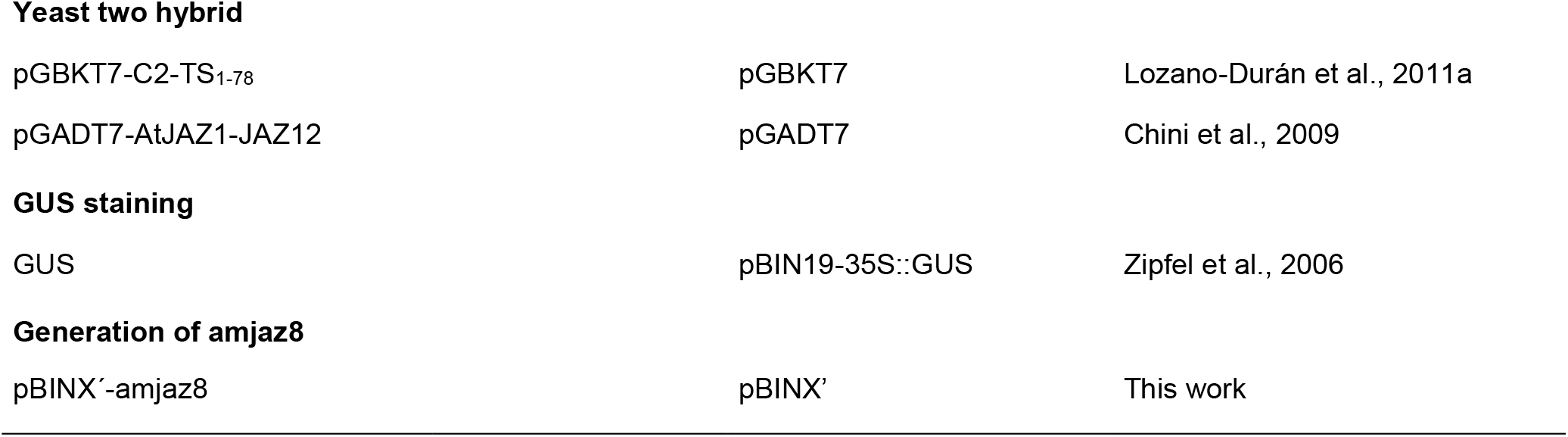
List of plasmids

**Supplementary Table 2.**
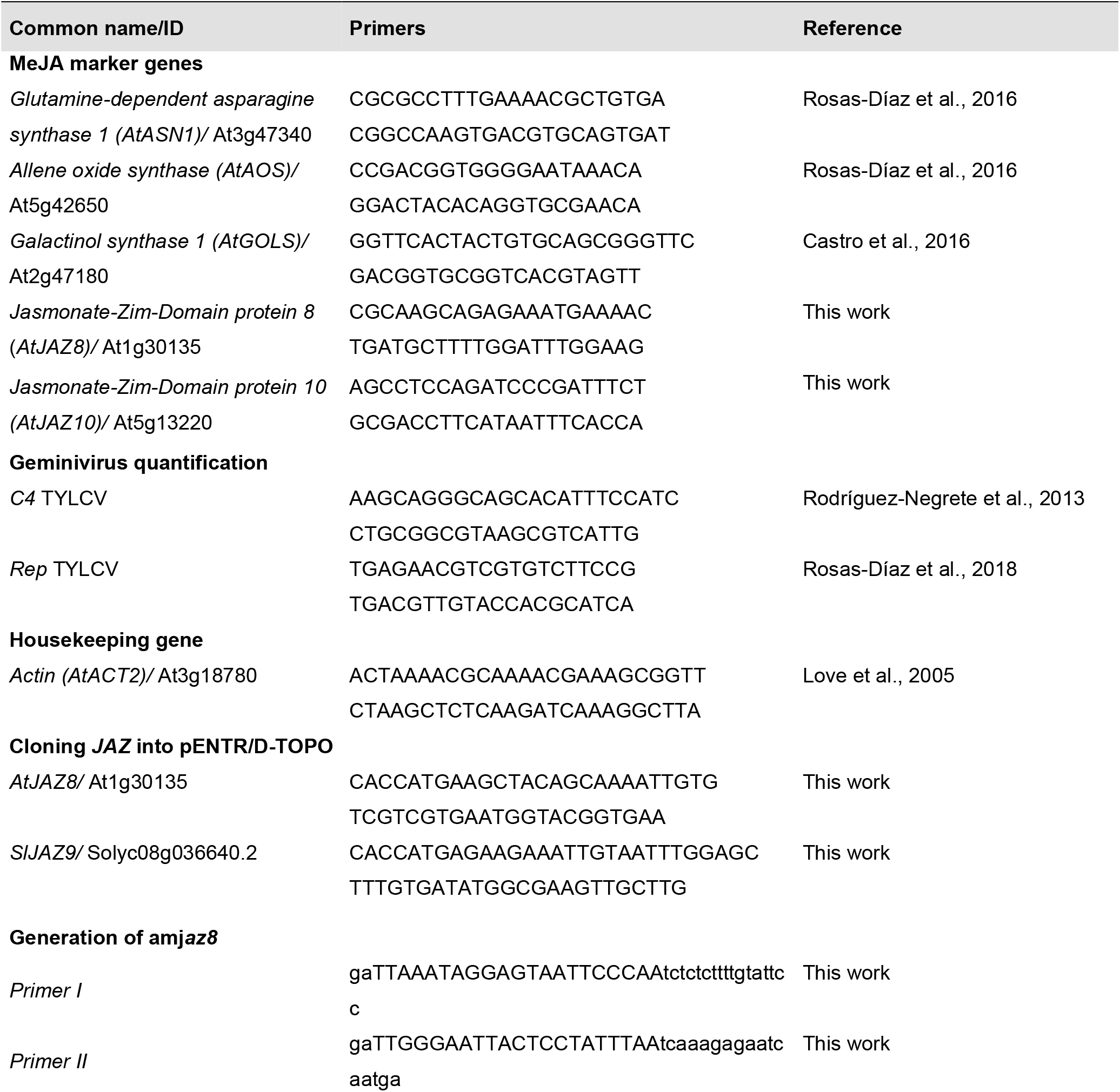
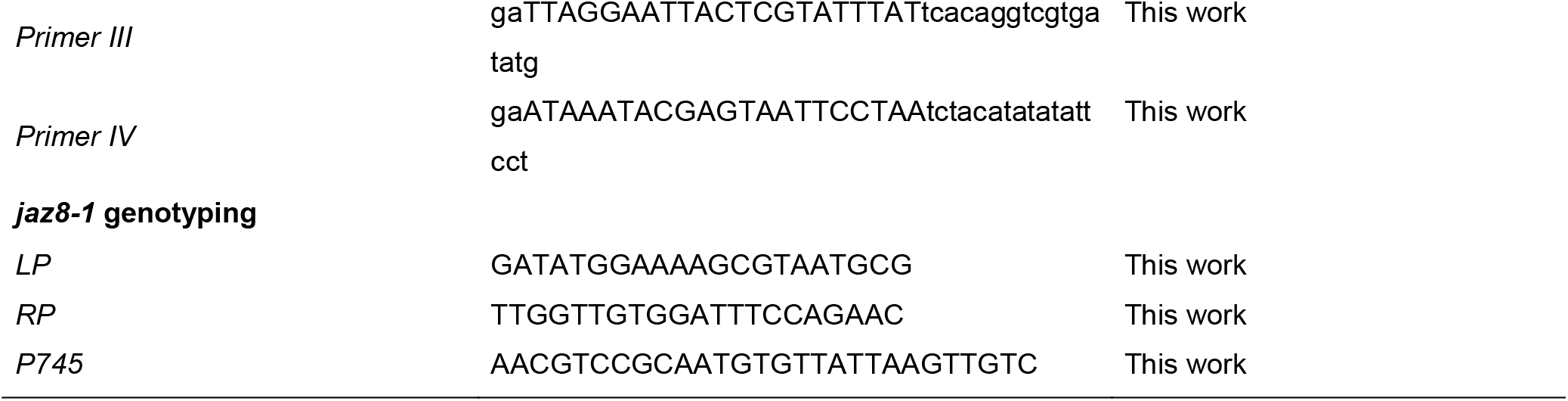
List of primers for PCR and qPCR

## Supplementary figures

**Supplementary figure 1.**
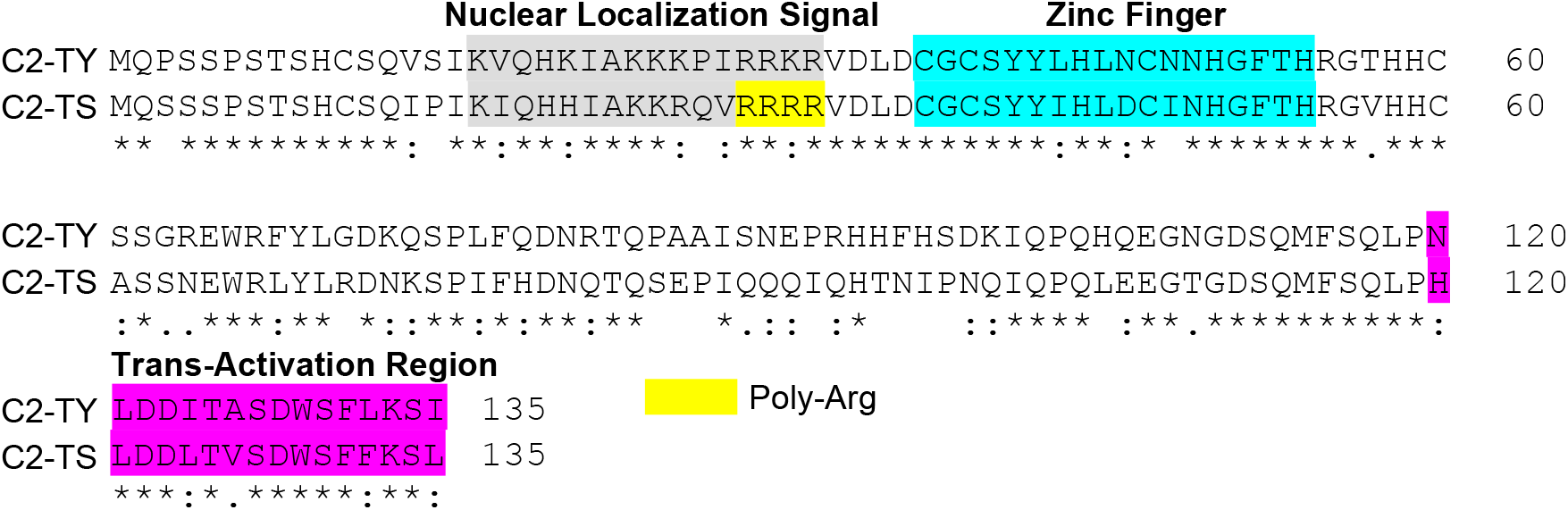
Alignment of C2 from begomoviruses TYLCV and TYLCSaV. The nuclear localization signal, zinc finger domain, and transactivation region are indicated. C2-TY: C2 from TYLCV (tomato yellow leaf curl virus) and C2-TS: C2 from TYLCSaV (tomato yellow leaf curl Sardinia virus); the sequence alignment was performed using Clustal Omega software (https://www.ebi.ac.uk/Tools/msa/clustalo/).

**Supplementary figure 2:**
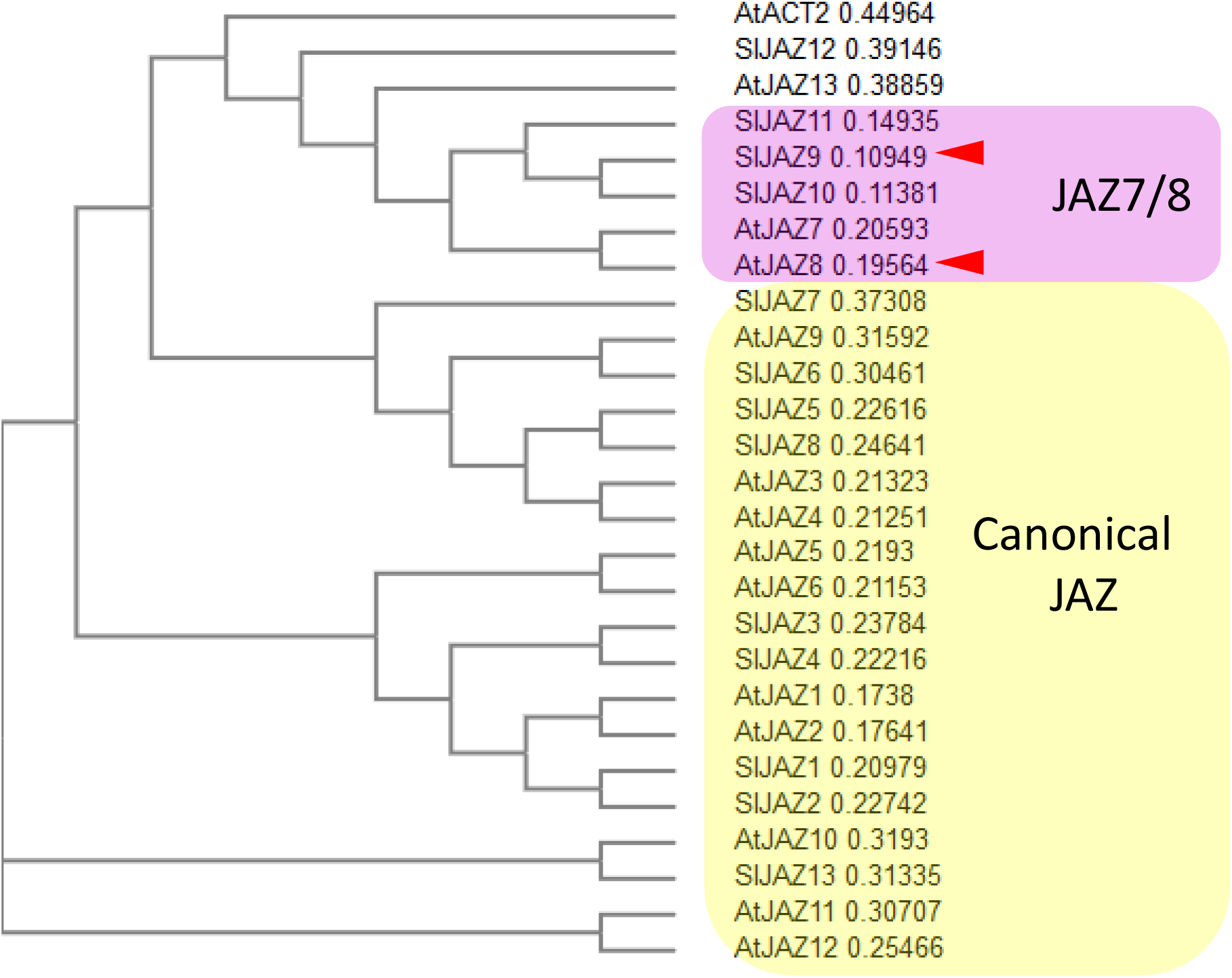
Phylogenetic tree of JAZ proteins. Phylogenetic analysis of *S. lycopersicum* and *A. thaliana* JAZ full-length proteins. The phylogenetic tree was created using the Clustal Omega software showing a Neighbour-joining tree without distance corrections (https://www.ebi.ac.uk/Tools/msa/clustalo/). Numbers correspond to the distance of branch length. Canonical JAZ proteins contain the canonical jas degron inside the jas motif.

**Supplementary figure 3.**
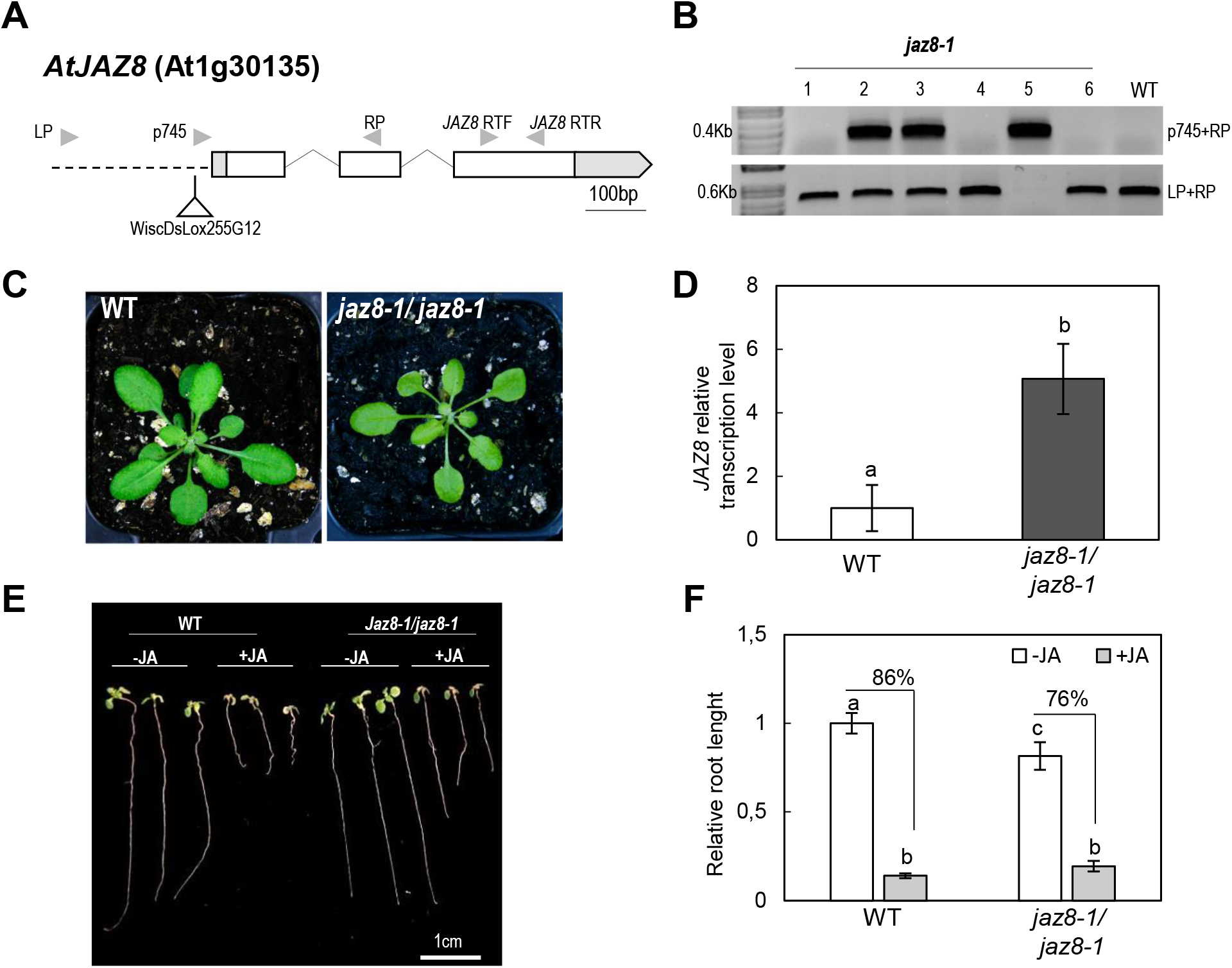
Characterization of the *JAZ8* T-DNA *Arabidopsis* line. **(A)** Diagram of *Arabidopsis JAZ8* gene showing the T-DNA insertion in *jaz8-1* line. Exons are represented in white boxes, introns as black lines and untranslated regions (UTR) as grey boxes. T-DNA insertion is located in the promoter region. The oligonucleotides used for PCR are depicted with arrows. **(B)** PCR products were obtained with P745+RP and LP+RP of six independent *JAZ8* T-DNA plants compared to wild type (WT). **(C)** The phenotype of three-week-old *Arabidopsis* plants compared to wild type plant (WT) and homozygous *jaz8-1* line.**(D)** Relative *JAZ8* mRNA levels in wild type, WT; and homozygous *jaz8-1* seedlings. Total RNA was extracted and subjected to quantitative RT-PCR (RT-qPCR) analysis to measure the *JAZ8* mRNA levels normalized to *ACTIN 2*. Values are represented as the relative expression compared to WT. Bars represent the mean +/-SD for three different pools of 8-10 seedlings. Similar results were obtained in two independent experiments. **(E)** Root growth inhibition assays in *jaz8-1* seedlings compared with *Arabidopsis* wild type (WT) and **(F)** relative root length of WT and *jaz8-1 Arabidopsis* seedlings growing at 50 μM MeJA (+JA) or mock (-JA) (0.5% ethanol in water (v/v)). Data has been relativized to WT (-JA) treatment values. In (D) and (F) mean values for at least ten seedlings are represented. Bars represent SE; bars with the same letters are not significantly different (P=0.05) according to Dunnett’s multiple comparison test. Numbers above bars indicates the percentage of root inhibition length upon JA treatment (+JA) compared to mock control (-JA) (100%). Experiments were repeated twice with similar results; results from one representative experiment are shown.

**Supplementary figure 4:**
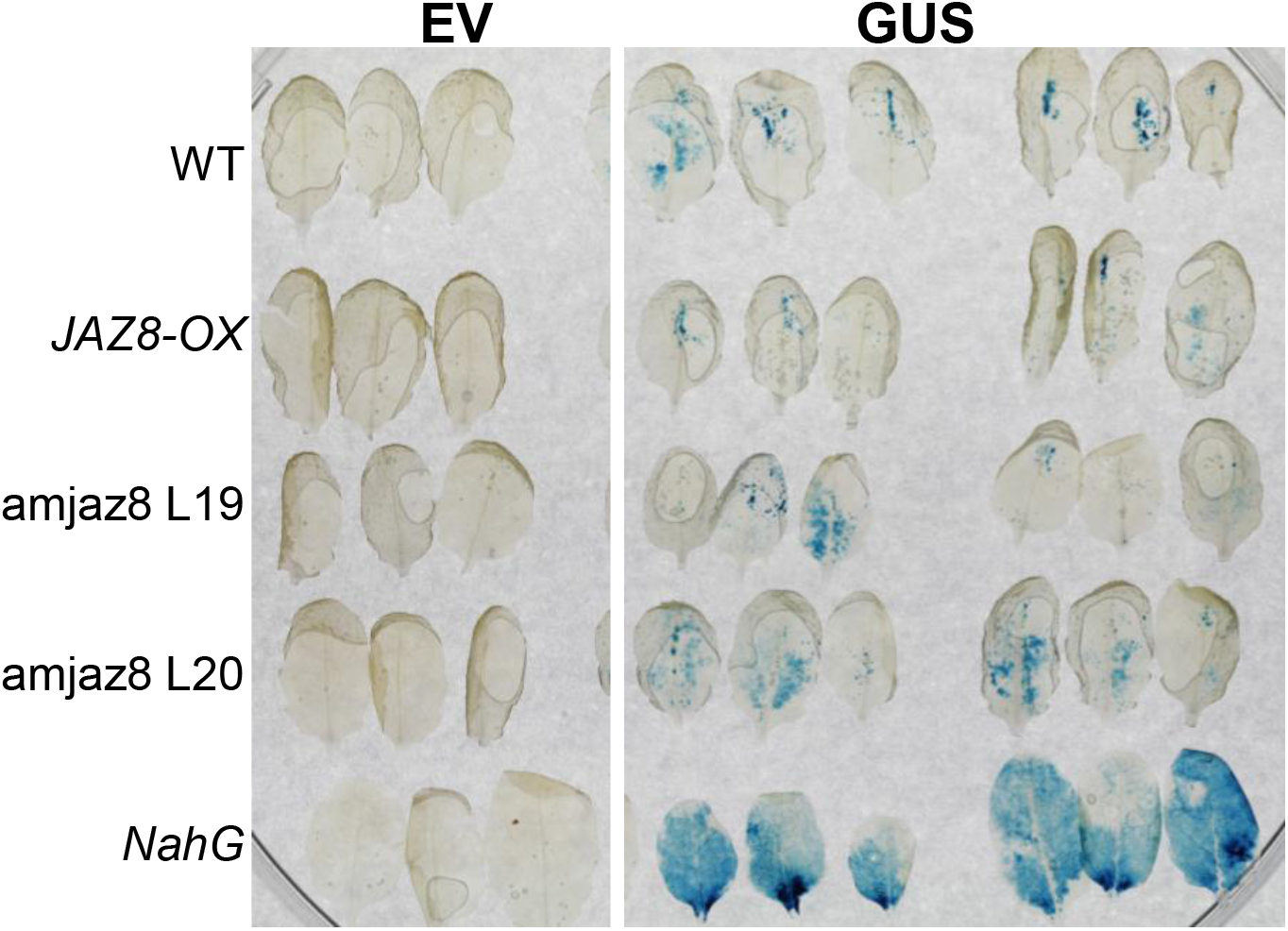
*Agrobacterium*-mediated T-DNA transfer is not affected by changes in *JAZ8* expression. Staining for GUS activity in leaves of wild type (WT), 35S:*JAZ8* (*JAZ8*-OX), amjaz8 (and *NahG* transgenic *Arabidopsis* plants after infiltration with 2.5*107 cfu/ml (DO600=0.05) *Agrobacterium* GV3101 carrying 35S::GUS or pBIN19 (EV) as a control. Photographs were taken at 4 dpi. Images are representative of 15 inoculated leaves per plant genotype.

## AUTHOR CONTRIBUTIONS

TR-D, PC-Q, GF-B, MW, AP-M and HD performed the experiments. RS, RL-D, ER-B have planned and directed the experimental design of the work. TR-D, RL-D and ER-B wrote the paper. AC-G participated in the discussion and experimental design of the work. All authors participated in the critical reading of the manuscript and contributed to the article and approved the submitted version.

## ACKNOWLEDGMENTS

This work was supported by the Spanish Ministerio de Ciencia y Tecnología (PID2019-107657RB-C22) (ER-B), the Shanghai Center for Plant Stress Biology, the Chinese Academy of Sciences, and the Federal Ministry of Education and Research (BMBF) and the Baden-Württemberg Ministry of Science as part of the Excellence Strategy of the German Federal and State Governments (RL-D). TR-D was supported by a President’s International Fellowship Initiative (PIFI) postdoctoral fellowship (No. 2016PB042) from the Chinese Academy of Sciences, the “Programa Juan de la Cierva” (IJCI-2017-33367) from the MCIN and FEDER program UMA20-FEDERJA-132 by AEI and by “ERDF A way of making Europe”, by the “European Union”. The authors would like to thank P. García Vallejo for his technical help.

